# Single-Cell Multiome Sequencing Clarifies Enteric Glial Cell Diversity and Identifies an Intraganglionic Population Poised for Neurogenesis

**DOI:** 10.1101/2021.08.24.457368

**Authors:** Richard A. Guyer, Rhian S. Stavely, Keiramarie Robertson, Sukhada Bhave, Ryo Hotta, Julia A. Kaltschmidt, Allan M. Goldstein

**Affiliations:** Department of Pediatric Surgery, Massachusetts General Hospital, Boston MA; Neurosciences Graduate Program, Stanford University, Stanford, CA; Department of Neurosurgery, Stanford University Medical Center, Stanford, CA

**Author notes:** Corresponding author: Allan M. Goldstein, MD Massachusetts General Hospital, 55 Fruit Street, Warren 1151, Boston, MA 02114.

**Keywords:** Enteric nervous system, enteric glial cells, glial cells, single-cell RNA sequencing, single-cell ATAC sequencing, single-cell multiome sequencing, neurogenesis

## Abstract

The enteric nervous system (ENS) consists of glial cells (EGCs) and neurons derived from neural crest precursors. EGCs retain capacity for large-scale neurogenesis in culture, and *in vivo* lineage tracing has identified neurons derived from glial cells in response to inflammation. We thus hypothesize that EGCs possess a chromatin structure poised for neurogenesis. We use single-cell multiome sequencing to assess EGCs undergoing spontaneous neurogenesis in culture, as well as freshly isolated small intestine myenteric plexus EGCs. Cultured EGCs maintain open chromatin at genomic loci accessible in neurons, and neurogenesis from EGCs involves dynamic chromatin rearrangements with a net decrease in accessible chromatin. Multiome analysis of freshly isolated EGCs reveals transcriptional diversity, with open chromatin at neuron-associated genomic elements. A subset of EGCs, highly enriched within the myenteric ganglia, has a gene expression program and chromatin state consistent with neurogenic potential.

## Introduction

The enteric nervous system (ENS) controls gastrointestinal (GI) motility, secretion, and absorption (Furness, 2012), and immune regulation (Jacobson et al., 2021). The ENS is composed of glial cells (EGCs) and neurons derived from migrating neural crest cells (ENCCs) during embryonic development (Nagy and Goldstein, 2017). Postnatal EGCs express many of the same markers as prenatal ENCCs, such as Sox10, p75, and Nestin (Joseph et al., 2011; Laranjeira et al., 2011), and EGCs have been demonstrated to generate neurons both in culture and *in vivo* in response to injury (Belkind-Gerson et al., 2015, 2017; Joseph et al., 2011). These data suggest EGCs maintain a multipotent state primed for neurogenesis.

Protein expression patterns of the glial markers Proteolipid Protein 1 (Plp1), Sox10, S100b, and Glial fibrillary acidic protein (Gfap) suggest heterogeneity among EGCs (Rao et al., 2015), but this has not been fully explored. Other evidence of functional diversity among EGCs include heterogeneous responses to purinergic stimulation, dye coupling of only a subset of glial cells within ganglia, and preferential stimulation of *Gfap* transcription in response to lipopolysaccharide exposure (Boesmans et al., 2015; Maudlej and Hanani, 1992; Rosenbaum et al., 2016). Several single-cell RNA sequencing (scRNA-seq) studies have captured EGCs and have reported multiple EGC clusters based on differential gene expression (Drokhlyansky et al., 2020; Zeisel et al., 2018). However, these scRNA-seq studies did not report detailed analysis of EGC subtypes, nor did they assess differentiation of EGCs into neurons. Our current work extends these results by jointly assessing transcription and chromatin structure of EGCs both in their native environment and during *in vitro* neurogenesis.

The gene expression programs available to a cell are constrained by epigenetic factors including chromatin accessibility (Wu et al., 2016). Open chromatin is sensitive to nuclease digestion and can be assayed by DNase I hypersensitivity or the assay for transposase-accessible chromatin (ATAC), the latter of which is a valuable tool for profiling chromatin in either bulk or single-cell samples (Boyle et al., 2008; Buenrostro et al., 2013; Satpathy et al., 2019). While differentiation is a dynamic process involving both closing and opening of chromatin (Jadhav et al., 2017), a net decrease in nuclease-sensitive regions accompanies progressive restriction in cell fate potential (Baumann et al., 2021; Stergachis et al., 2013; Ugarte et al., 2015). Similarly, perturbations that prevent terminal differentiation are associated with increased chromatin accessibility (Wenderski et al., 2020). Joint profiling of ATAC signal and gene expression can thus clarify how fate potential changes as cells adopt novel transcriptional programs.

We hypothesize that heterogeneous EGCs maintain a permissive chromatin structure for neuronal differentiation. In the present study, we apply scRNA-seq and single-cell multiome sequencing (scMulti-seq) to characterize EGCs and study the dynamics of neuronal differentiation. We show that EGCs maintain open chromatin at promoters and regulatory elements controlling neuronal marker genes. Differentiation of EGCs into neurons involves reduced chromatin accessibility, consistent with more limited fate potential. scMulti-seq of EGCs freshly isolated from postnatal mouse intestine reveals considerable transcriptional diversity, consistent with previous reports (Zeisel et al., 2018). By integrating gene expression with chromatin structure, we identify a subpopulation of EGCs primed for neurogenesis. *In situ* hybridization validates this population and demonstrates it is heavily enriched within myenteric ganglia. Our work provides a multiomic atlas of myenteric EGCs, identifies a chromatin state poised for neurogenesis, and shows intraganglionic EGCs to be biologically distinct from extraganglionic populations.

## Results

### Enteric Glial Cells Spontaneously Generate Neurons in Culture

We first sought to establish an *ex vivo* model for studying the EGC-to-neuron fate transition. We developed a dual-reporter system by crossing *Plp1::GFP* mice (Mallon et al., 2002), which mark all EGCs by GFP expression, with *Actl6b::Cre;*(*R)26-tdTomato* mice, which permanently mark cells committed to neuronal fate with tdTomato. We generated neurospheres using cells isolated from the longitudinal muscle-myenteric plexus (LMMP) layer of small intestine of these dual-reporter mice and then sorted GFP+/tdTomato-cells to obtain a pure EGC population. These EGCs were plated on fibronectin and live-cell imaging was performed 1 and 5 days later. The day after plating, no tdTomato+ cells were seen, indicating we successfully isolated a pure EGC population (Figure 1A). On day 5, however, a significant number of tdTomato+ neurons were noted, thus demonstrating that EGCs generate neurons in culture (Figure 1A). This confirms prior reports claiming EGCs have neurogenic potential in culture (Belkind-Gerson et al., 2015, 2017; Joseph et al., 2011; Laranjeira et al., 2011).

**Figure 1.**
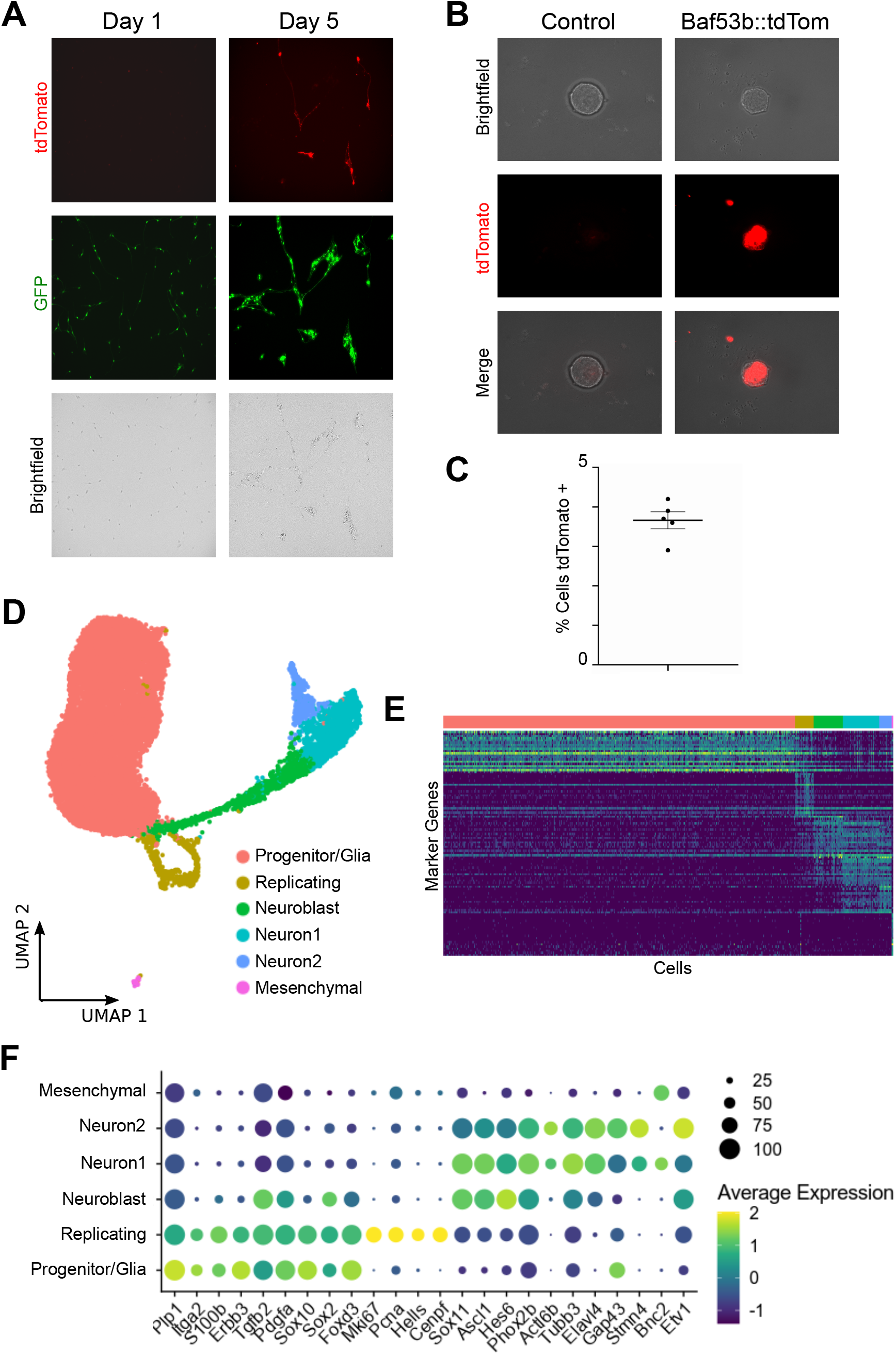
EGCs are neuronal progenitors in culture. A) GFP+/tdT-enteric glial cells isolated from *Plp1::GFP/ Baf53b::Cre;R26tdT* dual-reporter mice give rise to tdT+ neurons in culture. B) tdT-cells sorted from the small intestine of *Baf53b::Cre;R26tdT* mice generate tdT+ cells when grown as neurospheres. C) Flow cytometry quantification of tdT+ cells in neurospheres grown using tdT-cells sorted from the small intestine of *Baf53b::Cre;R26tdT* mice. D) Uniform Manifold Approximation and Projection (UMAP) of 15,426 GFP+ cells sorted from neurospheres grown from the LMMP cells of *Plp1::GFP* mice, with major cell types highlighted, demonstrates a continuum of gene expresssion patterns from glial cells to neurons. E) Single-cell heatmap with cells grouped by cluster, showing the top 25 marker genes for each cluster based on fold-change in expression relative to other clusters. F) Dot plot showing expression of selected marker genes for each cluster. Dot size indicates the percentage of cells in each cluster with >0 transcripts detected, while color indicates the relative level of gene expression.

To determine whether active neurogenesis occurs within neurospheres, we sorted tdTomato-negative cells from the small intestine LMMP of *Actl6b::Cre;*(*R)26-tdTomato* mice to obtain a population lacking committed neurons. We then generated neurospheres using this neuron-free population. After 7 days in neurosphere culture conditions, expression of tdTomato was apparent (Figure 1B), and flow cytometry analysis showed approximately 4% of cells within the neurospheres were tdTomato+ committed neurons (Figure 1C), demonstrating active neurogenesis occurring within neurospheres.

To better understand neurogenesis within neurospheres, we performed scRNA-seq on both the GFP+ and GFP-cell fractions sorted from neurospheres that were generated using LMMP cells from *Plp1::GFP* mice. The GFP-fraction was dominated by cells with a mesenchymal gene signature, which are known to promote ENS cell growth (Stavely et al., 2021). In contrast, the GFP+ fraction was heavily enriched for ENS marker genes (Figure S1A). As expected, the GFP-fraction included neurons (Figure S1B), which may represent neurons present at the time of gut dissociation or newly generated neurons in which GFP protein has been degraded. The GFP+ fraction includes a large number of EGCs, as expected, but also proliferating cells, cells expressing genes characteristic of enteric neuroblasts (Morarach et al., 2021), and neurons (Figures 1D and 1F). The neuroblasts and neuronal clusters contain progressively lower levels of Plp1 RNA (Figure 1F), and a cascade of gene expression changes from EGCs to neurons is evident (Figure 1E), which indicates neuronal differentiation from a glial cell of origin. The differentiation trajectory shows a similar pattern of gene expression changes as previously reported during embryonic ENS development in mice (Figures 1F, S1C, and S1D) (Morarach et al., 2021), including emergence of two distinct neuronal lineages marked respectively by *Bnc2* and *Etv1* transcription (Figures 1D, 1F, and S1D). Based on these data, we conclude that enteric neurosphere cultures are a useful model for studying enteric neurogenesis.

### Multiome Analysis of EGC-to-Neuron Transition Within Neurospheres Reveals a Chromatin Structure Poised for Neurogenesis

To evaluate whether EGCs maintain chromatin permissive for neurogenesis, we performed scMulti-seq to simultaneously obtain gene expression and ATAC data on GFP+ cells within neurospheres grown from *Plp1::GFP* mice. Dimensional reduction and clustering was performed using ATAC data, and cell identities were confirmed based on gene expression. This approach revealed a similar hierarchy as seen in our scRNA-seq data, with a large population of EGCs, two clusters of neurons, an intervening population of neuroblasts, and a cluster of proliferating cells (Figures 2A and 2B). To further confirm the validity of clustering via ATAC data, we used ChromVAR (Schep et al., 2017) to identify transcription factor binding motif enrichments. Consistent with a fate transition from glial cells to neurons, we found a switch from SOX10 and SOX2 motif enrichment to ASCL1 and PHOX2B motif enrichment (Figures 2C and S2A). As expected during differentiation from a multipotent progenitor to a more differentiated state, we observed a net closing of chromatin during the transition from EGCs to neurons (Figures 2D and 2E). The chromatin changes during the EGC-to-neuron transition are dynamic, with many EGC-associated ATAC peaks showing diminished signal in cell populations further along the differentiation vector. Conversely, chromatin becomes increasingly accessible at peaks associated with neuronal cells (Figure 2F). For example, several peaks around the Sox10 gene show markedly diminished signal as cells progress toward a neuronal fate (Figure 2H). Interestingly, many ATAC peaks that are associated with neuroblasts or neurons are nuclease-sensitive in EGCs (Figures 2F, 2G, and 2I), suggesting they are regulatory elements that can be activated to trigger neurogenesis.

**Figure 2.**
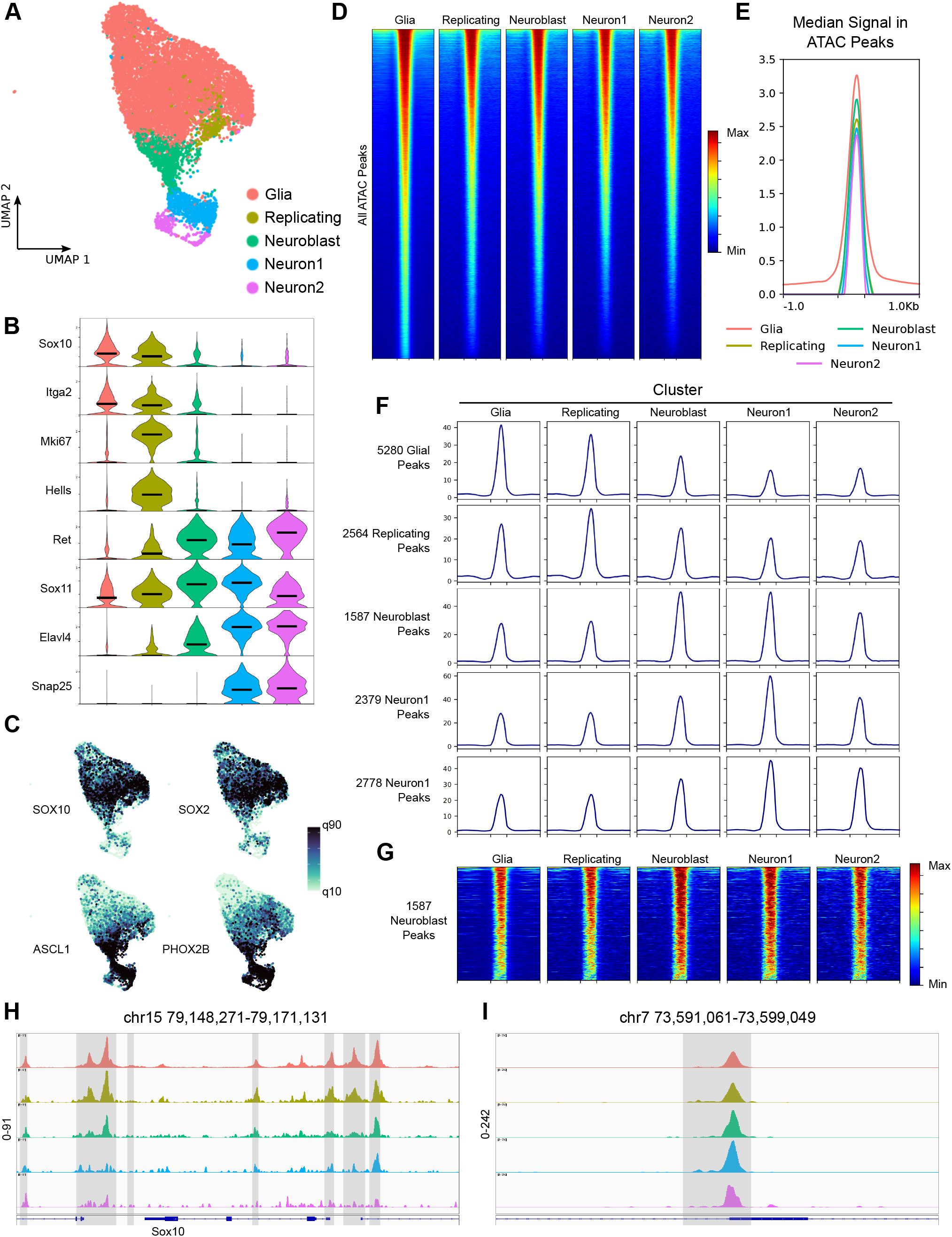
Multiome sequencing reveals chromatin poised for neurogenesis in EGCs in neurospheres. A) UMAP of 10,328 GFP+ cells sorted from neurospheres grown from the LMMP of *Plp1::GFP* mice, with dimensional reduction performed based on ATAC signal. Major cell types are highlighted. B) Violin plot showing expression of selected marker genes in each major cell type. Dark bars indicate median expression. C) UMAP projections colored based on enrichment for the indicated transcription factor binding motifs, with cutoffs at the 10^th^ and 90^th^ quantiles. D) Heatmaps showing signal within each cluster at all 171,799 ATAC peaks identified in the dataset by MACS2. Peaks are scaled to 500 bps, and 1000 bps up- and downstream are shown. E) Profile plot showing the median signal at each position within all 171,799 ATAC peaks. Peaks are scaled as in (D). F) Profile plots showing average signal intensity within each cluster at the indicated set of peaks. Peaks are scaled as in (D). G) Heatmaps showing signal at the indicated 1587 neuroblast peaks within each cluster. Peaks are scaled as in (D). H) IGV Browser track showing ATAC signal within each cluster in the region around the *Sox10* gene locus. Peaks identified by the MACS2 algorithm are highlighted in grey. I) IGV Browser track showing ATAC signal within each cluster in the region around an ATAC peak that marks neuroblasts. The peak region as identified by the MACS2 algorithm is highlighted in grey. The highest peaks are seen in the Neuroblast and Neuron1 clusters, but signal is also apparent in the Glia cluster.

We identified 578 genes whose RNA expression marks neuroblasts and neurons in our dataset. When individual cells were scored for a gene expression program defined by these genes, marked enrichment was seen in the neuroblast and neuron clusters (Figure S2B). We then identified ATAC peaks that are positively or negatively correlated with expression of these genes, and examined the ATAC signal within the coding regions of these genes, 2000 base pairs (bps) of upstream promoter, and 1000 bps downstream. To our surprise, the majority of upstream promoter regions for these 578 neuroblast- and neuron-associated genes were accessible in EGCs and displayed only a small increase in signal during neuronal differentiation (Figures S2C, S2D). This is illustrated by the neuronal marker gene *Tubb3*, which has very similar accessibility around the transcriptional start site (TSS) in EGCs, neuroblasts, and neurons (Figure S2E). The 578 genes were positively linked with 1634 ATAC peaks and negatively linked with 385 peaks. In contrast to signal in their promoter regions, there was a marked shift in signal at the linked peaks (Figures S2B and S2C), with dramatic shifts in nuclease sensitivity along the trajectory of neurogenesis. We conclude that the promoter regions of neuronal lineage-defining genes are primed for rapid activation in EGCs, and can be rapidly induced when regulatory elements are activated or repressed.

### EGCs in the Postnatal Small Intestine are Transcriptionally Diverse and Include Cells With a Chromatin Structure Primed for Neurogenesis

Although neurospheres derived from LMMP cells can model enteric neurogenesis, it is unclear whether EGCs in these cultures reflect the biology of EGCs *in vivo*. We thus undertook scRNA-seq and scMulti-seq of GFP+ cells freshly isolated from the small intestine LMMP of *Plp1::GFP* mice at around postnatal day 14 (P14). We integrated these datasets and performed dimensional reduction and clustering based on gene expression. We filtered out low-quality cells and manually removed small numbers of cells that co-expressed markers of both glial cells (*Sox10, Plp1*) and either neurons (*Tubb3, Uchl1*) or smooth muscle cells (*Acta2*). Although prior literature has suggested that EGCs can generate both neurons and myofibroblasts (Joseph et al., 2011; Zeisel et al., 2018), we could not rule out the possibility of doublet contamination, so these cells were excluded from further analysis. We were left with 17,690 EGCs, 3754 of which had both gene expression and ATAC data. Clustering based on differential gene expression identified 9 clusters of EGCs (Figure 3A). We observed two clusters of proliferating cells marked by cell cycle-associated genes (*Mki67, Cenpf, Hells, Pcna*). Interestingly, we noted an inverse correlation between expression of several glial markers, including *Plp1, Sox10, Sox6*, and *Itga2* (which were high in clusters 0, 2, 3, 7, and 8), and *Gfap, S100b*, and *Sox2* (high in clusters 1 and 4, Figure 3B). Clusters 1 and 4 were marked by high expression of *Slc18a2, Ramp1*, and *Cpe* (Figure 2B), the first of which was previously identified as a marker of a subset of EGCs in mouse colon (Drokhlyansky et al., 2020). Clusters 1 and 4 were also relatively enriched for genes encoding transcription factors associated with neuronal differentiation, including *Phox2b, Hand2, Tbx3, Ascl1, Hoxa5*, and *Hoxb5* (Figure 3B). We also found that genes previously identified as EGC markers in bulk RNA sequencing studies, such as *Kcna1* and *Col20a1* (Rao et al., 2015), had low expression in clusters 1 and 4 relative to other EGCs (Figure 3B). These data show that postnatal EGCs are a heterogeneous population that remains proliferative and includes a subset (clusters 1 and 4) expressing genes associated with neurogenesis.

**Figure 3.**
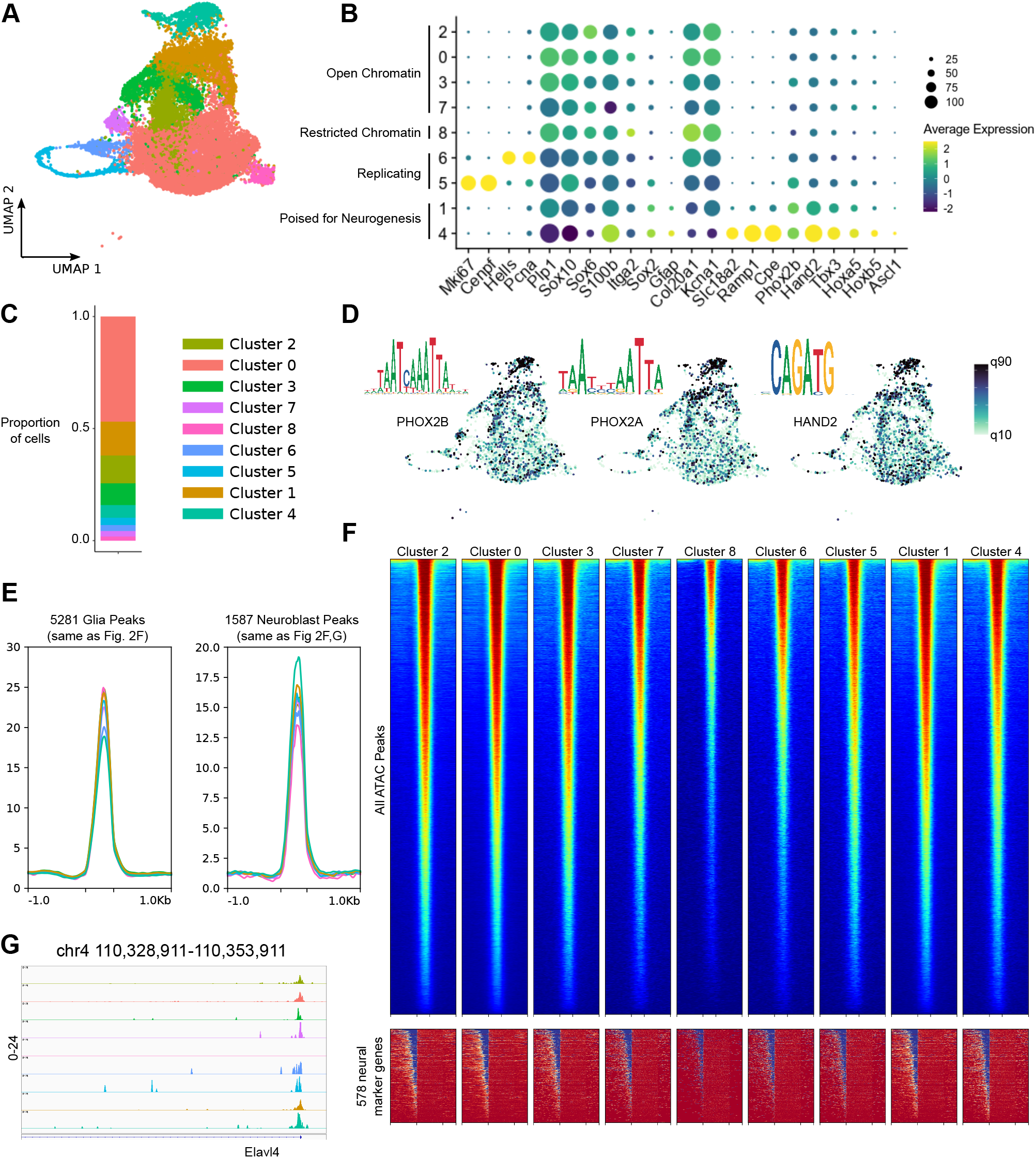
Small intestinal myenteric glia are transcriptionally diverse and contain cells poised for neurogenesis. A) UMAP of 17,690 GFP+ cells sorted from the small intestine of *Plp1::GFP* mice near P14, with dimensional reduction performed based on differential gene expression. B) Dot plot showing expression of selected genes within cluster. Dot size indicates the percentage of cells in each cluster with >0 transcripts detected, while color indicates the relative level of gene expression. Clusters are annotated to the left based on gene expression, chromatin accessibility at neuronal marker peaks, and motif enrichment patterns. C) Proportion of cells contained within each cluster. D) UMAP projections colored based on enrichment for the indicated transcription factor binding motifs, with cutoffs at the 10^th^ and 90^th^ quantiles. Also shown are position frequency plots for the indicated motifs. E) Profile plots showing average signal intensity at the indicated set of peaks. Peaks are scaled to 500 bps, and 1000 bps up- and down-stream are shown. F) Heatmaps showing ATAC signal within each cluster at all 94,210 ATAC peaks identified in the dataset by MACS2 (top), and around the gene body of 578 marker genes for neuroblasts and neurons identified within the dataset (bottom). ATAC peaks (top) are scaled to 500 bps, and 1000 bps up- and downstream are shown, while gene bodies (bottom) are scaled to 2000 bps, and 2000 bps up- and 1000 bps downstream are shown. G) GV Browser track showing ATAC signal within each cluster in the region around the *Elavl4* TSS.

For further confirmation that clusters 1 and 4 contain EGCs poised for neurogenesis, we again used ChromVAR. Among the motifs enriched in clusters 1 and 4 we noted PHOX2B, PHOX2A, and HAND2, all of which are factors associated with neuronal differentiation (Morarach et al., 2021). We examined ATAC signal at the 5481 EGC marker peaks and 1588 neuroblast marker peaks identified in our neurosphere model (Figure 2F). We noted that ATAC signal at the EGC peaks was diminished in cluster 4, while ATAC signal was enriched at neuroblast peaks in clusters 1 and 4 (Figure 3E). We also noted that cluster 8 showed the strongest inverse correlation with cluster 4 in both peak sets. This suggests EGCs in clusters 1 and 4 have a chromatin structure permissive for neurogenesis, while cluster 8 may represent EGCs that have lost the capacity to become neurons.

We next examined global chromatin accessibility. We found most glial clusters to display a similar degree of chromatin accessibility, with somewhat diminished signal in proliferating cells. The notable exception was cluster 8, which had a significantly lower ATAC signal at most peaks (Figure 3F). We assessed ATAC signal in the promoters and coding regions of the 578 neuronal marker genes previously identified (Figures S2B). As in the neurospheres, most EGCs *in vivo* maintain accessible chromatin upstream of these genes’ TSS. Cluster 8 was again noted to be an exception, with significantly diminished signal relative to other clusters (Figure 3F). This is illustrated by the neuronal marker Elavl4, which shows similar ATAC signal around its TSS in all clusters except cluster 8 (Figure 3G). These results indicate that EGCs in the postnatal mouse intestine contain a subpopulation primed for neurogenesis (clusters 1 and 4), as well as another subpopulation (cluster 8) with a restricted chromatin structure and decreased chromatin accessibility near the TSS of neuronal marker genes.

### In Situ Hybridization Validates EGC Diversity and Reveals EGCs Poised for Neurogenesis are Restricted to Ganglia

We used RNAscope (Wang et al., 2012) to confirm that clusters 1 and 4 represent a genuine subpopulation of EGCs. LMMP from the small intestine of *Plp1::GFP* mice was used for this analysis. Because Gfap protein expression has been reported to mark a subset of EGCs (Rao et al., 2015), we first evaluated *Gfap* transcripts. Consistent with the scRNA-seq data, *Gfap* was detected in only a subset of EGCs (Figures 4A and S3A), although the proportion of EGCs with *Gfap* transcripts detected was considerably higher by *in situ* hybridization. This discrepancy may reflect greater sensitivity of RNAscope for detection of genes with low transcript abundance. Consistent with Rao et al. (2015), we observed that *Gfap* transcripts are considerably enriched within the myenteric ganglia as compared to extraganglionic EGCs (Figures 4A and S3A). Transcripts of four other genes that are co-enriched with *Gfap* in clusters 1 and 4 were also evaluated (*Sox2, Cpe, Ramp1*, and *Slc18a2*). In each case, enrichment of these transcripts within myenteric ganglia was apparent (Figure 4A). Quantification of the staining showed that cells dual-positive for *Gfap* transcripts and each of these four transcripts represented a significant majority of cells within the myenteric ganglia (Figures 4B-4E). Transcripts of *Slc18a2* were particularly scarce outside of the ganglia (Figure 4E). In addition to being detected in a smaller number of cells outside of ganglia, the mean flouresence intensity (MFI) of the signal from these four genes was lower in extraganglionic EGCs (Figures S3B-S3E), indicating lower transcript levels. The lone exception was *Cpe*, which showed no difference in MFI between intragangionic and extraganglionic Ggap+ cells (Figure S3C), although intraganglionic Gfap+ cells did have significantly higher MFI of the *Cpe* probe than either intra- or extraganglionic Gfap-negative cells (Figure S3C). These results validate the presence of an EGC subpopulation marked by coexpression of *Gfap, Sox2, Cpe, Ramp1*, and *Slc18a2* transcripts, which represents cells apparently poised for neurogenesis, and demonstrates that these cells are heavily enriched within myenteric ganglia.

**Figure 4.**
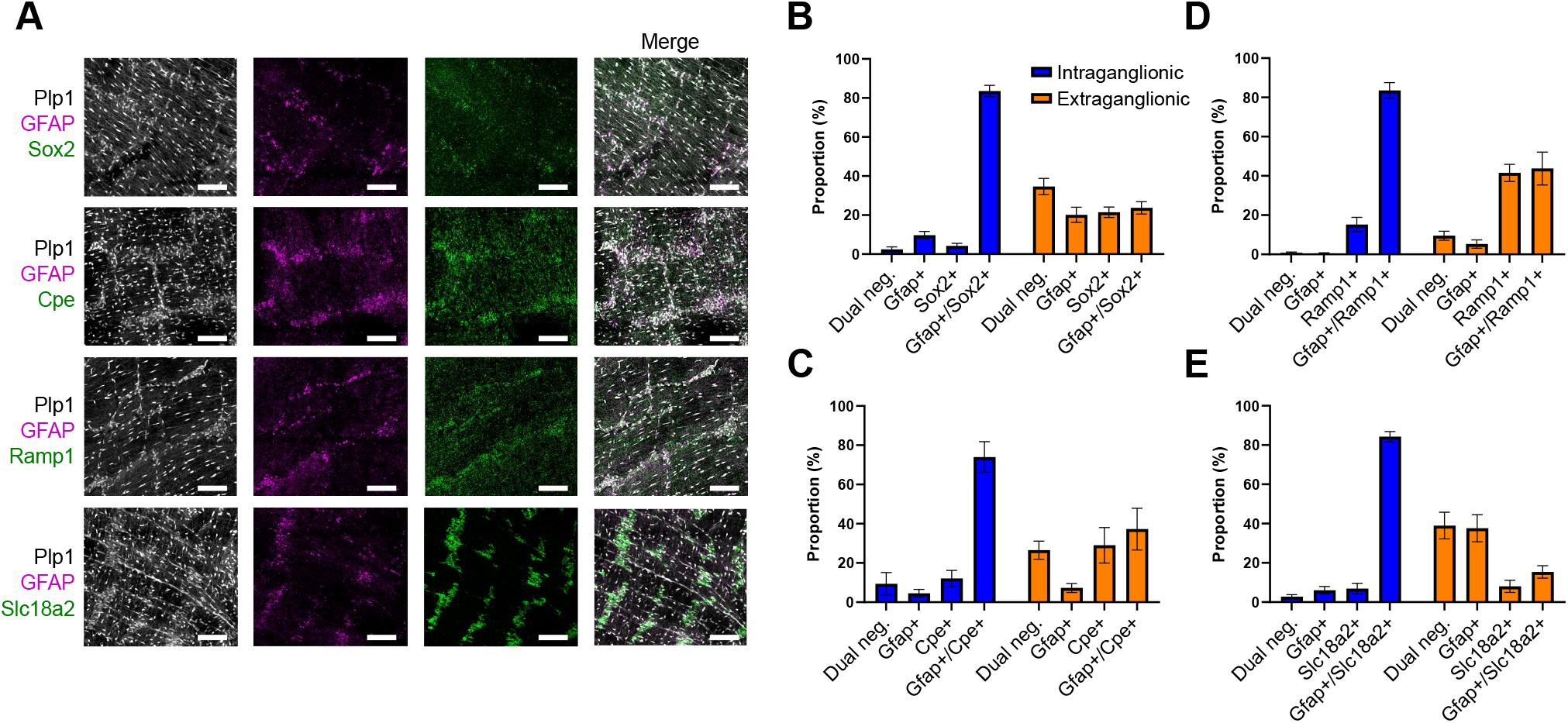
RNAscope confirms an EGC subpopulation marked by *Gfap* transcription and localizes these cells to myenteric ganglia. A) Confocal microscopy images of RNAscope staining of EGCs in the LMMP of Plp1::GFP mice around P14, with staining for *Gfap* (magenta) and the other genes indicated (green). Scale bar indicates 100 mm. B-E) Quantification of the proportion of intra- and extraganglionic EGCs with RNAscope signal for the indicated combinations of genes.

## Discussion

Postnatal EGCs are a heterogeneous population of cells (Rao et al., 2015; Zeisel et al., 2018) shown to possess the capacity to generate neurons both in culture and *in vivo* in response to injury (Belkind-Gerson et al., 2015, 2017; Joseph et al., 2011; Laranjeira et al., 2011). Our present work, by using multiome sequencing technology to examine cultured and *in vivo* EGCs, confirms that EGCs are progenitors of enteric neurons, identifies intraganglionc glial cells as having a transcriptional and epigenetic state poised for neurogenesis, and identifies a chromatin state in EGCs that is likely to maintain multipotency. In addition, we demonstrate that neurogenesis in neurosphere culture closely recapitulates the transcriptional events observed during embryonic enteric neuronal development (Morarach et al., 2021). Since neurospheres are a tractable experimental model and can be generated in large numbers from a single mouse intestine, they represent an excellent system for studying the molecular mechanisms of enteric neurogenesis. The similarity between glial-derived neurogenesis in neurosphere cultures and embryonic neurogenesis further suggests that large-scale expansion of EGCs as neurospheres *in vitro* may be a valuable source of enteric neuronal progenitors for regenerative cell therapies, a promising area of active investigation for the treatment of neurointestinal diseases (Burns et al., 2016).

By utilizing scMulti-seq, we have clarified the molecular basis underlying the neuronal potential of enteric glia. We find that EGCs have a more open chromatin structure than enteric neurons, which is a hallmark of cells at earlier points along a differentiation vector (Baumann et al., 2021; Stergachis et al., 2013; Ugarte et al., 2015). As EGCs become neurons, their chromatin changes dynamically, with closing of EGC-associated sites and opening at neuronal-associated loci both occurring. EGCs, both in culture and in their native environment, maintain open chromatin at loci that characterize neuroblasts and neurons. The set of accessible chromatin sites likely includes regulatory elements that can be activated to initiate a cascade of transcriptional and epigenetic changes that initiates a lineage commitment and culminates in a neuronal fate transition. Such a process has been demonstrated in the mouse hair follicle, where increased ATAC signal at enhancers and promoters precedes changes in gene expression that determine cell fate (Ma et al., 2020). Further characterization of this process will require profiling DNA modifications, histone marks, and transcription factor binding during neurogenesis, which will undoubtedly be facilitated by the emergence of tools for these types of single-cell analysis (Ahn et al., 2021; Bartosovic et al., 2021; Kaya-Okur et al., 2019).

Interestingly, we have found that EGCs both in culture and in the small intestine LMMP maintain open chromatin upstream of the transcription start sites of many neuronal marker genes despite low expression of these genes. ATAC signal is generally correlated with gene expression (Wu et al., 2016), although cells can maintain accessible chromatin within the promoters of genes that are not actively transcribed (Starks et al., 2019) and ATAC signal within promoters is a sign of impending transcription during development (Ma et al., 2020). High ATAC signal within neuronal marker genes’ promoters in EGCs thus suggests poising for neurogenesis. We speculate that EGCs repress neuronal lineage-determining genes via mechanisms other than chromatin accessibility to prevent inappropriate neurogenesis, and that such repression is relieved upon receipt of differentiation signals. Given these observations, EGCs may display bivalent chromatin within promoters of neuronal lineage-determining genes, which would allow for their rapid activation (Bernstein et al., 2006), but this remains to be determined.

We have used *in situ* hybridization with RNAscope to validate transcriptional heterogeneity among myenteric EGCs. RNAscope identified *Gfap* transcripts in a considerably higher proportion of EGCs than scRNA-seq, a discrepancy likely due to greater sensitivity of the RNAscope assay, although inefficient isolation of single cells from ganglia may also contribute. Nevertheless, the results confirm that *Gfap* transcription correlates with *Sox2, Cpe, Ramp1*, and *Slc18a2. Slc18a2* has previously been identified as an EGC cluster marker in the mouse colon and small intestine (Drokhlyansky et al., 2020; Zeisel et al., 2018), which demonstrates that scRNA-seq experiments are robust for identifying EGC populations. *Gfap*-expressing EGCs, which comprise clusters 1 and 4 in our scRNA-seq data, also express RNAs that encode transcription factors associated with neurogenesis (such as *Phox2b, Phox2a, Ascl1*, and *Hoxa5*), have higher signal at neuron- and neuroblast-associated ATAC peaks than other EGC clusters, and are enriched for neuronal transcription factor binding motifs. Based on these data, we believe *Gfap*-expressing EGCs represent glial cells poised for neurogenesis. Interestingly, RNAscope revealed that the *Gfap*-expressing EGCs are heavily enriched within the myenteric ganglia. This recapitulates the results of Rao et al. (2015), who found via immunostaining that Gfap protein is largely confined to ganglia. We speculate that spatial restriction reflects different functions for intra- and extraganglionic EGCs. Intraganglionic EGCs may be a reservoir of potential neurons, while the greater expression of extracellular matrix genes, such as *Col20a1*, and ion channel genes, such as *Kcna1*, in extraganglionic EGCs may reflect neuronal support roles and other functions important for intestinal homeostasis.

In summary, this study provides a multiome atlas of mouse EGCs and provides a valuable, publicly available resource for the scientific community. By integrating gene expression and ATAC data at the single-cell level, we have found that the transition from EGC fate to neuronal fate involves dynamic epigenome rearrangements. A subset of EGCs within the myenteric plexus that is heavily enriched within the myenteric ganglia has a gene expression program and chromatin state poised for neurogenesis. These data will inform studies to determine critical signaling networks and transcriptional circuitries underlying the EGCs’ decision to either remain multipotent or become a neuron.

### Limitations of the Study

The postnatal mice in this study are adolescent pups around P14. Changes to glial cells during early adulthood and later in life are beyond the scope of this report but may be important, and our data cannot assess whether EGCs fate potential is altered in older animals. Our study is limited to EGCs from the LMMP, so neither submucosal EGCs nor Schwann cells accompanying extrinsic nerve fibers are included. Extrinsic Schwann cells have been suggested as a source of myenteric neurons in adult animals and may have similar epigenetic potential (Uesaka et al., 2015). Our data reflect a pooled population of EGCs from the entire small intestine, and colonic cells are not included. Regional variation in chromatin state or gene expression, as previously reported (Drokhlyansky et al., 2020; Rao et al., 2015), could be masked. Although ATAC data allows for inference of regulatory elements and transcription factor binding patterns (Akerberg et al., 2019; Galang et al., 2020), profiling of histone modifications and transcription factor binding can add additional information, but is outside the scope of this paper. Finally, as is the current standard, we have used enzymatic digestion and cell sorting to isolate EGCs from the LMMP layer. Such techniques can be confounded by artifactual gene signatures in diverse tissues, including brain microglia (Adam et al., 2017; Marsh et al., 2022). Multiome sequencing including ATAC can mitigate this problem, as ATAC signal delineates cell identity more robustly than RNA (Corces et al., 2016).

## Acknowledgments

The authors are grateful for expertise offered by the Harvard University Bauer Core and the Harvard Stem Cell Institute Center for Regenerative Medicine Flow Cytometry Core facility. This work was supported by the National Institute of Diabetes and Digestive and Kidney Diseases (grants F32DK121440 to RAG and R01DK119210 to AMG), by the Department of Neurosurgery of Stanford University School of Medicine (JAK), by the Firmenich Next Generation Fund (JAK). KR’s work was supported by National Institute’s of Health training grant T32MH020016 awarded to Stanford University School of Medicine.

## Author Contributions

Conceptualization – RAG and AMG; Methodology – RAG, AMG, KR, JK, and RSS; Formal Analysis – RAG and RSS; Investigation – RAG, SB, and KR; Resources – AMG, RH, and JK; Data Curation – RAG and RSS; Writing of Original Draft – RAG and AMG; Review and Editing – RAG, RSS, KR, JK, and AMG; Supervision – AMG, RH, and JK; Project Administration – AMG; Funding Acquisition – RAG, AMG, and JK.

## Methods

### Animals

This study was performed according to experimental protocols approved by the Institutional Animal Care and Use Committees of Massachusetts General Hospital and Stanford University. *Plp1::GFP* mice were kindly gifted to the Goldstein laboratory by Wendy Macklin (Mallon et al., 2002) or purchased from Jackson Laboratories (Bar Harbor, ME) (stock number 033357). Animals homozygous for GFP expression were used for scRNA-seq, while animals heterozygous for GFP expression were used for RNAscope studies. *Actl6b::Cre* (stock number 027826) and (*R)26-tdTomato* (stock number 007914) mice were purchased from Jackson Laboratories (Madisen et al., 2010; Zou et al., 2015). Dual reporter mice were generated by first crossing *Actl6b::Cre* animals with (*R)26-tdTomato* animals, and subsequently crossing offspring with *Plp1::GFP* animals to yield *Plp1::GFPI/ Actl6b::Cre;*(*R)26-tdTomato* offspring.

### Isolation of ENS Cells

Mice of the indicated ages and genotypes were euthanized and their small intestine was removed from duodenum to terminal ileum. The longitudinal muscle-myenteric plexus (LMMP) layer, which contains myenteric ganglia, was carefully dissected from underlying tissue under a dissecting microscope in ice-cold PBS supplemented with 10% bovine serum albumin. After dissection, LMMP tissue was digested for 60 minutes at 37° C in dispase (250 μg/ml; STEMCELL Technologies, Vancouver, BC) and g/ml; STEMCELL Technologies, Vancouver, BC) and collagenase XI (1mg/ml; Sigma-Aldrich, St. Louis, MO). Following digestion, the cells were filtered via a 40 micron filter to ensure a single-cell suspension and either taken immediately for sorting or placed in neurosphere culture.

### Cell Sorting

Cell sorting was performed by the Harvard Stem Cell Institute’s Center for Regenerative Medicine Flow Cytometry Core facility located on the Massachusetts General Hospital campus. Sorting was performed on BD Bioscences (Franklin Lakes, NJ) FACSAria sorting instruments.

### Neurosphere Cultures

Immediately after digestion and filtering, cells were counted and resuspended at a density of 10^5^ cells/ mL in a 1:1 mixture of DMEM (Thermo Fisher, Waltham, MA) and NeuroCult Basal Media (STEMCELL Technologies, Vancouver, BC) supplemented with 20 ng/mL FGF, 20 ng/mL IGF1, 2% B27 supplement, 1% N2 supplement, 50 mM b-mercaptoethanol, and 75 ng/mL retinoic acid. 10^5^ cells/ mL in a total volume of 10 mL were placed 10cm flasks (Corning Inc, Corning, NY) at 37° C and 5% CO_2_ for 10 days. Media was replaced on day 5 by centrifuging cells at 250g for 3 minutes followed by resuspension in the same media and return to the same flasks and incubator for 5 more days. Mice age 12-16 weeks were used to obtain cells for neurosphere cultures.

### Preparation of scRNA-seq Libraries

For postnatal mice, LMMP cells were isolated from *Plp1::GFP* mice and sorted for GFP+ cells as described above. Immediately after sorting, cells were manually counted with Trypan blue to assess viability. Cells were then delivered immediately to the MGH NextGen Sequencing Core Facility and core staff prepared cDNA libraries using the 10X Genetics v3.0 kits. The standard 10X Genetics workflow was used. Cells from 4 mice were used in three batches, with the first batch containing pooled cells from 1 male and 1 female mouse and the remaining two batches each containing cells from a single mouse (1 male, 1 female). For scRNA-seq of three-dimensional (3D) cultures, single cells were obtained by digesting cultures for 45 minutes with Accutase (STEMCELL Technologies, Vancouver, BC). Cells were then sorted as described above into GFP+ and GFP-populations and immediately returned to the laboratory where they were manually counted with Trypan blue to assess viability. After determining viable cell counts, a 10X Chromium Controller located in our facility was used along with 10X Genetics (Pleasanton, CA) v3.1 kits to generate gel bead emulsions (GEMs), followed by library preparation according to the manufacturer’s protocol. Cells from 4 mice were used to generate 3D cultures for scRNA-seq, with cells from 2 male mice and 2 female mice each pooled together and cultured separately.

### Preparation of snATAC and Multiomic Libraries

For adolescent mice, LMMP was isolated from P16-18 animals as described above and GFP+ cells were sorted. Cells from three male mice were pooled and run on one lane of a GEM chip. 10X Genetics Multiomic kits were used, and GEMs were generated using the 10X Chromium Controller located in our facility. snATAC libraries were generated using the standard 10X Genetics workflow. For multiomic analysis of neurosphere cultures, LMMP cells from three 12-week-old *Plp1::GFP* mice were pool and grown in suspension culture conditions for 10 days. Cells were then dissociated and sorted to isolate the GFP+ population. A single lane on a GEM chip was then used to generate GEMs. The 10X Genetics multiomic library preparation workflow was undertaken in accordance with the manufacturer’s protocol.

### Sequencing and Genome Alignment

All sequencing was performed at the Harvard University Bauer Core Facility, where libraries were sequenced on either Illumina NextSeq or Illumina NovaSeq instruments. Demultiplexing, genome alignment, and feature-barcode matrix generation was performed with the 10X Genetics Cell Ranger software pipeline (Zheng et al., 2017).

### Single Cell Data Analysis

scRNA-seq and snMulti-seq data was analyzed with the open-source Seurat and Signac packages implemented in the R computing environment. For the postnatal glial scRNA-seq dataset and neurosphere datasets, cells more than one standard deviation away from the mean number of genes detected were filtered, as were cells with greater than 10% mitochondrial genes. Datasets were integrated using the SCTransform workflow in Seurat (Hafemeister and Satija, 2019). After integration, principle component analysis (PCA) was performed. Neighbors were identified and UMAP projection was performed using the first 30 principal components. Clusters were identified using the “FindClusters” command with resolution = 0.5 using the Louvain algorithm (Blondel et al., 2008). Where indicated, data were manually annotated based on expression of known marker genes.

ATAC data was processed using the standard Signac workflow. Briefly, cells with fewer than 1000 or more than 100,000 ATAC fragments were filtered, as were cells with nucleosomal enrichment > 2 or transcriptional start site enrichment < 1. Peaks within each dataset were identified using MACS2 (Zhang et al., 2008). Dimensional reduction was performed with latent semantic indexing (LSI) via the “RunTFIDF” command, “FindTopFeatures” function with min.cutoff set to 5, and “RunSVD” function. UMAP projection was performed utilizing LSI components 2-50.

Transcription factor motif enrichement was implemented with the ChromVAR software package implemented through Signac (Schep et al., 2017). JASPAR 2020 vertebrate transcription factor motifs were utilized (Fornes et al., 2020). ChromVAR results were imported to Signac as an assay object. To identify a gene set characteristic of neuroblasts and neurons, gene markers for each clusters were identified using the Seurat “FindAllMakers” command with the following settings: min.pct = 0.25, test.use = “wilcox”, only.pos = TRUE, logfc.threshold = 0.25. Unique marker genes of the clusters “Neuroblast,” “Neuron1,” and “Neuron2” were then selected and filtered to include only those with an adjusted p value < 0.0001 and a log2-fold change > 0.5 relative to other clusters. ATAC peaks linked with gene expression were calculated by using the Signac “LinkPeaks” command with default settings. Peak sets and gene loci were exported in BED format (Kent et al., 2002).

### ATAC Data Visualization

BAM files for each cluster were generated using Sinto (https://timoast.github.io/sinto/). Cluster BAM files were used to generate bigWig files using the deepTools (Ramírez et al., 2014) bamCoverage function with RPGC normalization, binSize =10, smoothLength = 50, effectiveGenomeSize = 2652783500, and extendReads = 150. deepTools computeMatrix was then used with the scaling indicated for each figure, and plots were generated using deepTools plotHeatmap or plotProfile. For heatmaps, zMin was set to 0 and zMax was set to the 90^th^ percentile value for the cluster with the highest median expression, as determined by using the deepTools computeMatrixOperations dataRange command. Tracks plots were generated using the Integrative Genomics Viewer (Robinson et al., 2011).

### Live Cell Imaging

GFP+/tdTomato-cells were sorted from LMMP suspension cultures derived from *Plp1::GFPI/ Actl6b::Cre;*(*R)26-tdTomato* dual reporter mice. Cells were placed in adherent culture conditions in neuronal differentiation media, which consists of BrainPhys Neuronal Media (STEMCELL Technologies, Vancouver, BC) supplemented with 1% N2 supplement and 2% NeuroCult S1 supplement (STEMCELL Technologies, Vancouver, BC). After one day media was removed and replaced with a thin layer of phosphate buffered saline. Cells were then imaged at 10x magnification using a Keyence microscope. After imaging, differentiation media was replaced and cells were returned to culture for 4 more days. Repeat imaging was performed in the same manner after 5 days in culture.

### RNAScope

Mice were euthanized and their small intestine was removed from the duodenum to the ileum. The small intestine was cut into three segments and each segment was cut along the mesentery. The cut segments were laid flat on filter paper and fixed for 24 hours in 4% PFA. Under a dissecting microscope, the LMMP was carefully peeled away from the lamina propria. Protein-RNA co-detection was performed using Advanced Cell Diagnostics (ACD) RNA-protein Co-detection ancillary kit (323180) according to manufactures instructions with adaptions for LMMP sections. Briefly, LMMP sections were post fixed in 4% PFA for 15 minutes in a 12-well plate. Tissue was transferred into staining nets and dehydrated by a serial ethanol gradient (50%, 70%, 100%, 100%) for 5 minutes each. Tissue was then placed into a 96-well plate and incubated in hydrogen peroxide for 15 minutes. Tissue was briefly rinsed in water and incubated in co-detect target antigen retrieval solution for 5 minutes in a steamer. Tissue was rinsed in water and incubated overnight with anti-GFP antibody (Abcam, ab13970) diluted 1:250 in co-detection diluent. RNAscope was performed using ACD RNAscope multiplex fluorescent reagent kit V2 (323100). All RNAscope incubation and washes were performed in 80-well microtitration trays (International Scientific Supplies, WHO080). Briefly, tissue was washed in 0.2% PBT and post-fixed in 10% NBF for 30 minutes at room temperature. Tissue was digested with Protease Plus for 25 minutes at 40°C. Tissue was incubated for 2 hours in RNAscope probes for *Gfap* (313211-C3), *Sox2* (401041-C2), *Slc18a2* (425331), *Ramp1* (532681-C2) or *Cpe* (454091). Negative and positive control probes were used to confirm specificity of probes and presence of background noise. Amplification and probe development steps were performed according to manufacture instruction. Tissue was incubated for 30 minutes in Alexa Flour 647 Goat anti-chicken secondary antibody (Invitrogen, A21449) diluted at 1:250 in co-diluent. Tissue was then mounted on slides and cover slipped for imaging. A Leica SP8 confocal microscope was used to acquire large tile images. Tiles were stitched together using the Navigation mode in the LASX software. Z-stacks with 2µm between each focal plane were acquired for 25-35µm thick sections. A 20x oil objective was used to acquire all images.

Quantification of RNAscope labeling was performed using Fiji (ImageJ v1.53c, NIH, USA) using methods previously described for the measurement of fluorescence in confocal images (Schindelin et al., 2012). Glial cells were identified by *Plp1* promoter-driven expression of GFP in Z-stack projections. Binary thresholding was performed on GFP expression using the default ImageJ algorithm. Glial cell bodies were identified using the “analyze particles” feature for objects over 20µm^2^. Individual GFP-expressing glia were manually annotated based on intra- or extraganglionic location, and 100 intraganglionic and 100 extraganglionic cells were arbitrarily selected from each sample for further analysis. ROIs generated from the particle analysis of individual glia were used to measure the background corrected mean fluorescence intensity (MFI) (Shihan et al., 2021) for RNAscope probes in each cell. Cells with a MFI >1 were considered to be positively labeled for calculating the proportion of cells expressing each marker.

### Published scRNA-seq Data

Data of the mouse embryonic ENS at P15.5 and P185 was published previously (Morarach et al., 2021). This data was obtained from the NCBI Sequence Read Archive. Runs SRR11635571, SRR11635572, and SRR11635573 were downloaded using the SRA Toolkit “fastq-dump” command. Genome alignment and feature-barcode matrix generation was performed with the Cell Ranger “cellranger count” command on the Mass General Brigham ERISOne Research Computing Cluster. Further analysis was performed with Seurat in the R environment. Briefly, the datasets were filtered as the authors describe in the original paper, with removal of cells with < 1000 genes detected, > 6000 genes detected, > 40,000 total RNA counts, and > 5% mitochondrial genes detected. Datasets were then integrated using the standard Seurat workflow. Following integration, PCA was performed and UMAP projection was undertaken with the top 30 principle components.

## Figure Legends

**Supplemental Figure 1.**
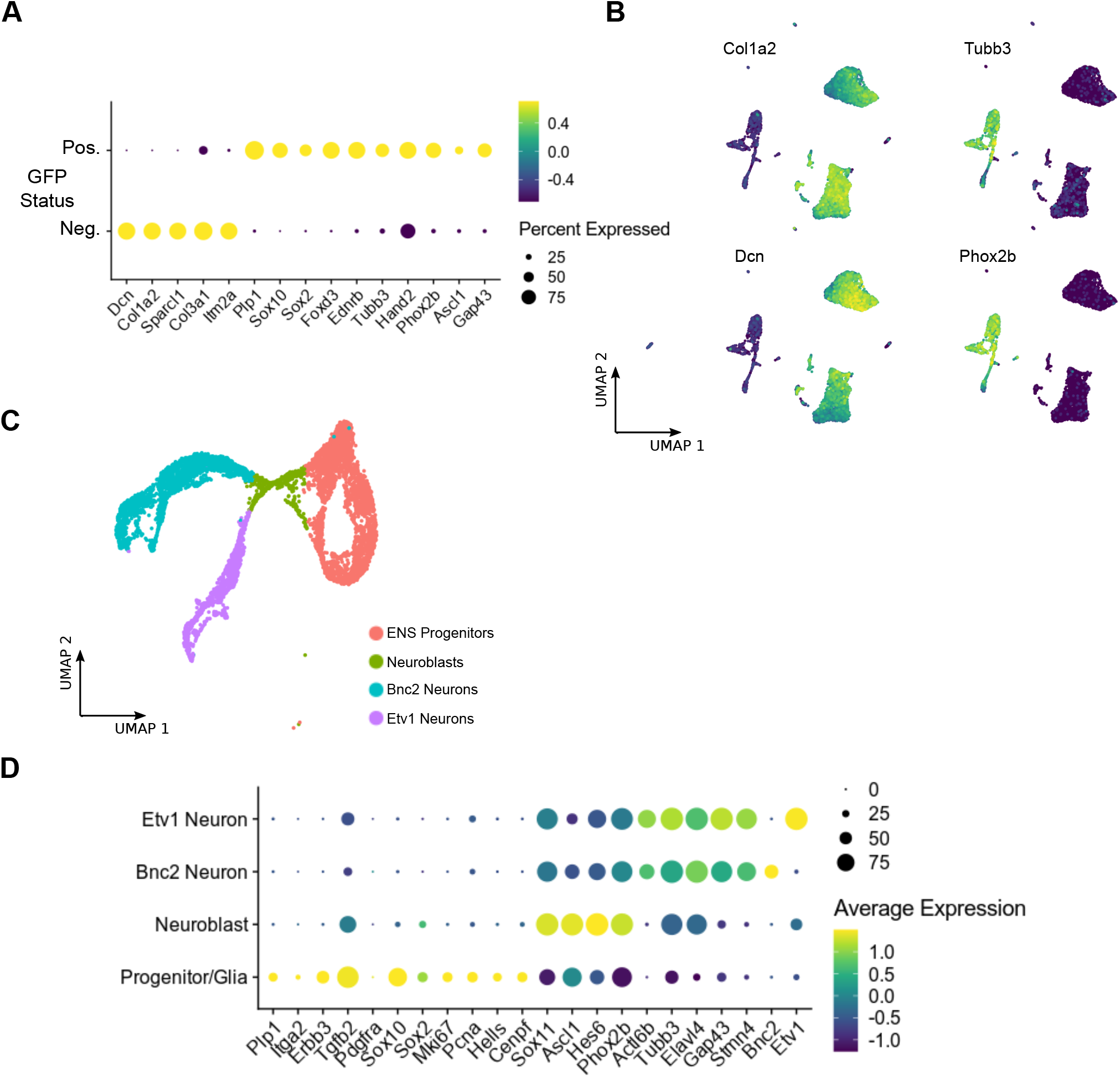
(related to Figure 1) A) Dot plot showing expression of selected genes in the GFP+ and GFP-negative populations of cells isolated from neurospheres grown from the LMMP of *Plp1::GFP* mice. Dot size indicates the percentage of cells in each cluster with >0 transcripts detected, while color indicates the relative level of gene expression. B) UMAP projections showing transcript abundance for the indicated genes within single cells of the GFP-population. C) UMAP projection showing the combined E15.5 and E18.5 murine embryonic ENS datasets, previously published by Morarach et al. (2021). Cells were manually assigned to the groups indicated based on gene expression. D) Dot plot showing expression of the same genes as in Figure 1F for each cluster in the combined embryonic dataset shown in (C). Dot size indicates the percentage of cells in each cluster with >0 transcripts detected, while color indicates the relative level of gene expression.

**Supplemental Figure 2.**
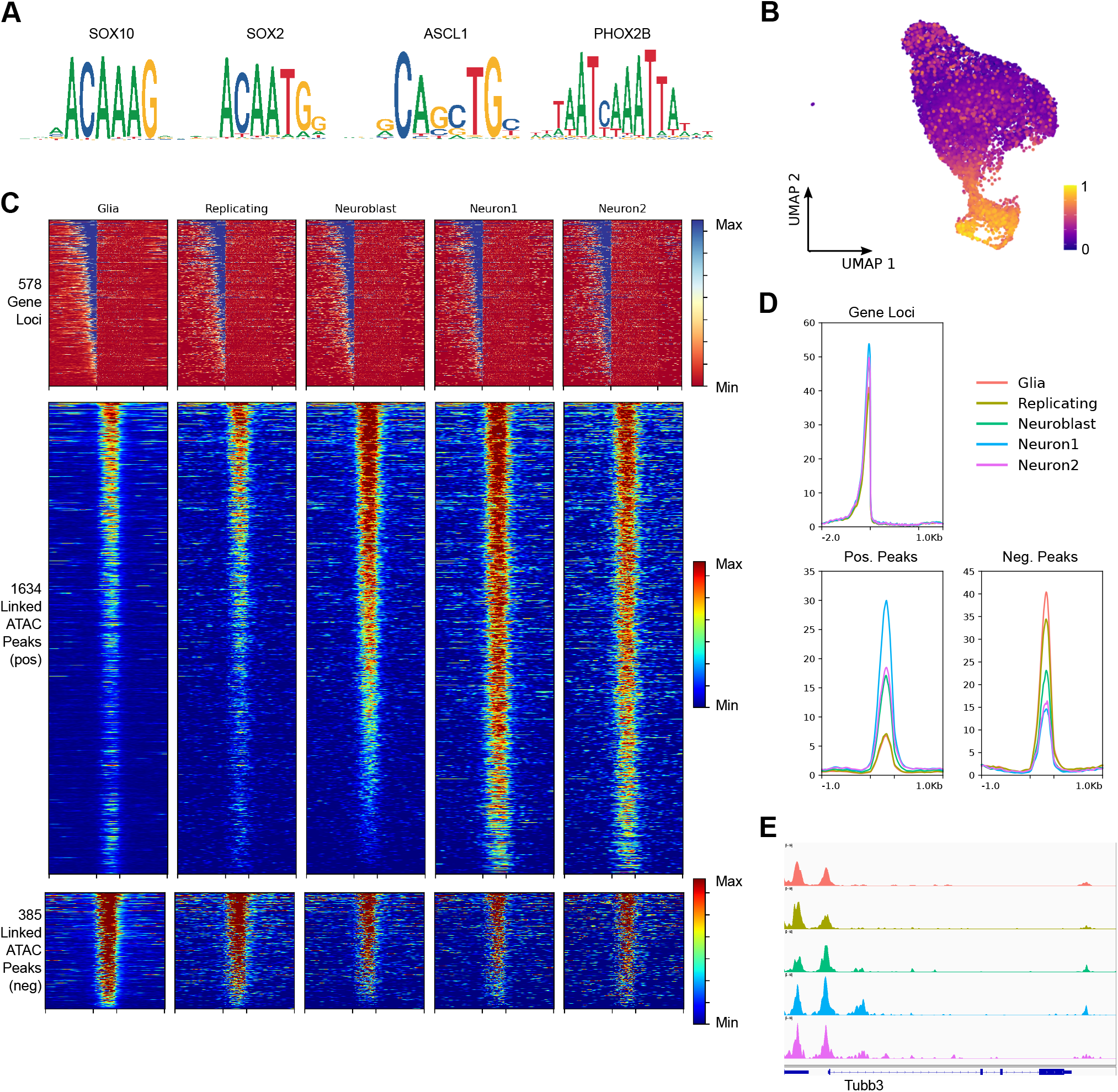
(related to Figure 2) A) Position frequency plots for the transcription factor binding motifs indicated, enrichment for which is shown in Figure 2C. B) UMAP projection showing the neuronal gene module score in individual cells. The module score was calculated based on expression of 578 neuroblast and neuronal marker genes. C) Heatmaps showing ATAC signal around the gene body of 578 marker genes for neuroblasts and neurons (top), 1634 ATAC peaks whose signal positive correlates with expression of the 578 marker genes (middle), and 385 ATAC peaks whose signal negatively correlates with expression of the 578 marker genes (bottom). Gene bodies (top) are scaled to 2000 bps, with 2000 bps upstream and 1000 bps downstream included, while ATAC peaks (middle and bottom) are scaled to 500 bps, with 1000 bps up- and down-stream included. D) Profile plots showing average ATAC signal around 578 neuroblast and neuronal marker gene loci, 1634 positive-correlated ATAC peaks, and 385 negatively-correlated ATAC peaks, as indicated. Regions are scaled as in (C). E) IGV Browser track showing ATAC signal within each cluster in the region around the *Tubb3* gene locus.

**Supplemental Figure 3.**
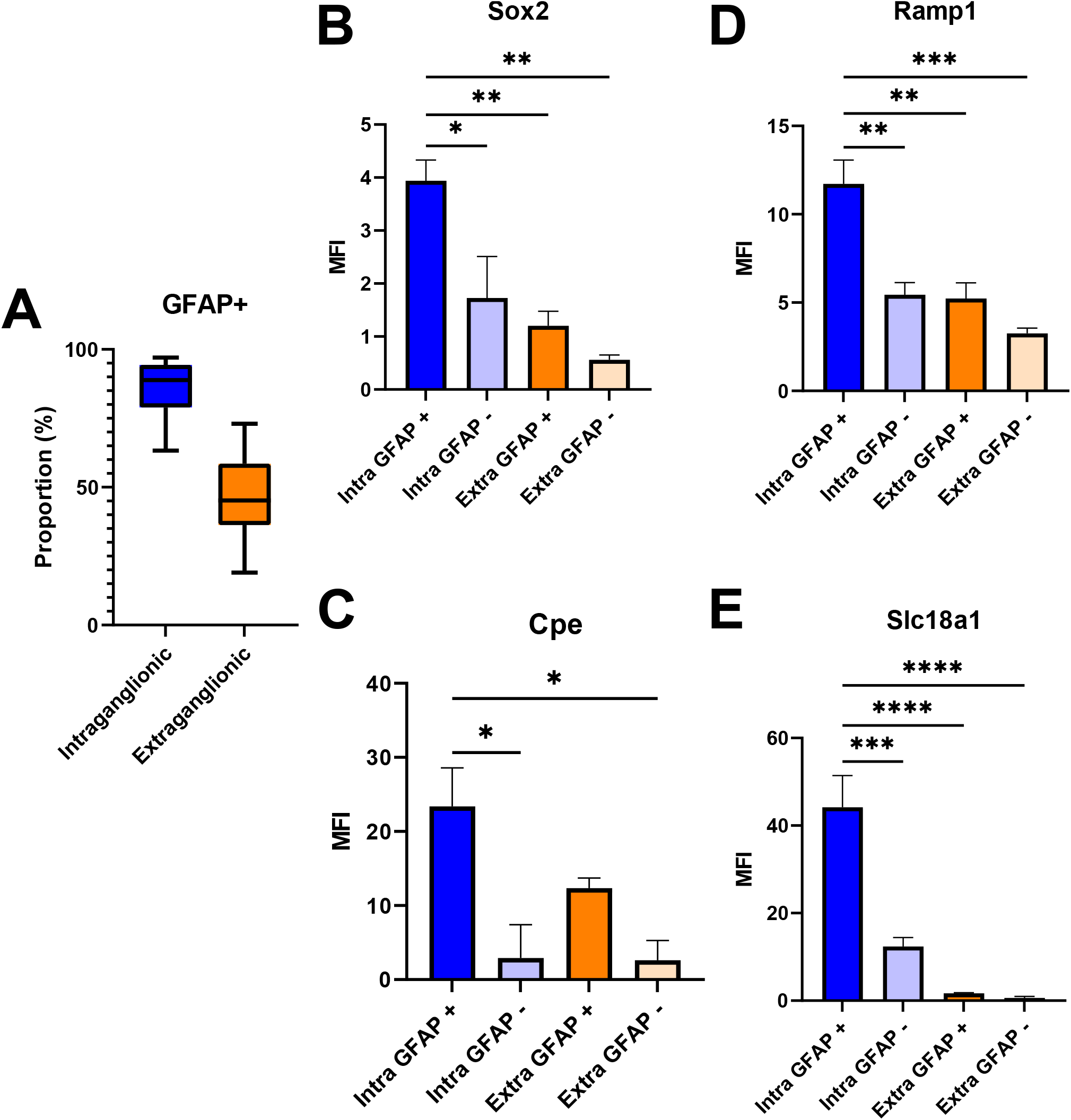
(related to Figure 4) A) Boxplot showing the proportion of intra- and extraganglionic cells in which *Gfap* transcripts are detected. Horizontal dark line indicates median, boxes indicate the interquartile range, and thin bars indicate minimum and maximum. p < 0.0001 for intraganglionic vs extraganglionic. B-E) MFI for the probes detecting the indicated transcripts, stratified by intra- or extraganglionic location and *Gfap* expression. *, p < 0.01; **, p < 0.001; ***, p < 0.0001; ****, p < 0.00001.

